# Bedaquiline inhibits the ATP synthase leak channel and prevents glutamate-induced neuronal death

**DOI:** 10.64898/2026.01.05.697836

**Authors:** Amrendra Kumar, Erin Smith, Ikram Mezghani, Subhash Eedarapalli, Yangyu Wu, Khondoker Adeba Ferdous, Emma Amjad, Han-A Park, Nelli Mnatsakanyan

**Affiliations:** Department of Cell and Biological Systems, Pennsylvania State University College of Medicine, Hershey, PA, USA; Department of Cellular and Molecular Physiology, Yale University, New Haven, CT, USA; Department of Human Nutrition, Hospitality and Sport Management, University of Alabama, Tuscaloosa, AL, USA

## Abstract

F_O_F_1_-ATP synthase is one of the most abundant proteins of the mitochondrial inner membrane and the primary enzyme responsible for ATP production in eukaryotic cells. Nevertheless, it was recently reported to play a prominent role in cell death by forming a large-conductance leak channel in the mitochondrial permeability transition pore (mPTP), making it a promising therapeutic target. Bedaquiline (BDQ), a member of the diarylquinoline class of drugs, was shown to selectively inhibit the catalytic activity of *Mycobacterium tuberculosis* ATP synthase with no effect on the mammalian enzyme. Here, we report a new role for BDQ as a potent inhibitor of the ATP synthase c-subunit leak channel in mammals. BDQ inhibited the single-channel activity of porcine heart ATP synthase in planar lipid bilayer recordings and prevented glutamate-induced cell death in primary hippocampal neurons. These findings reveal the potential new application of BDQ for treating mPTP-related diseases by targeting the ATP synthase c-subunit leak channel.

**Why it matters:** Bedaquiline (BDQ) is the only FDA-approved drug to treat pulmonary multidrug-resistant tuberculosis (TB), caused by the *Mycobacterium tuberculosis*. BDQ cures TB by specifically targeting mycobacterial ATP synthase and inhibiting ATP production. Recently, BDQ was also reported to bind to mammalian ATP synthase at the interface between the a and c-subunits and to inhibit its catalytic activity. However, the effect of BDQ on ATP synthase leak channel activity has not been explored. Here, we report that BDQ inhibits the ATP synthase c-subunit leak channel (ACLC) activity with an IC50 of ∼24 nM and prevents glutamate-induced neuronal death, suggesting a new therapeutic repurposing of BDQ for treating ACLC-related diseases.

## Introduction

The mitochondrial permeability transition pore (mPTP) plays a central role in cell death during ischemia-reperfusion (I/R) injury of the heart and brain, and in neurodegenerative diseases^1, 2^. The prolonged opening of mPTP leads to mitochondrial swelling and rupture, the release of cytochrome c and other mitochondrial signaling molecules, and the activation of cell death via apoptotic or necrotic pathways^2, 3^. Changes in inner mitochondrial membrane (IMM) permeability transition (mPT) occur due to the activation of the large-conductance, non-selective megachannel of the IMM, of unknown molecular composition. Over the years, various mitochondrial proteins, such as the voltage-dependent anion channel (VDAC)^4, 5^, the phosphate carrier (PiC)^6^, the translocator protein (TSPO)^7^, and the adenine nucleotide translocator (ANT)^8^, have been suggested as core components of the mitochondrial permeability transition pore (mPTP). However, studies involving the genetic deletion of these proteins have shown that they function more as modulators of the pore rather than as its structural or pore-forming elements^9-12^.

Over the past decade, growing evidence suggests a role of ATP synthase in mPTP formation^2, 13-16^. Purified mammalian ATP synthase forms a large-conductance, voltage-gated ion channel in electrophysiological recordings, with a similar peak conductance to that of mPTP and sensitivity to divalent and trivalent ions (Ca^2+^, Ba^2+^, Gd^3+^)^14, 17, 18^, and cyclophilin D (CypD)^19^. According to one of the current models, the channel is located between the ATP synthase dimers^13,18^. F_O_ subunits e and g that facilitate ATP synthase dimerization were shown to be involved in channel formation^16, 20-22^. In addition, purified ATP synthase c-ring was reported to form a voltagegated channel, and its genetic ablation eliminated the Ca^2+^-sensitive large-conductance activity of mitochondria, supporting a possible role of c-ring in mPTP formation^23, 24^. A low conductance channel activity, sensitive to ANT inhibitor bongkrekic acid (BA), was still present in c-subunit knockout cells, suggesting the possible contribution of multiple pores to mPT^23, 25^. Liver mitochondria from mice with triple deletion of *Ant1, Ant2*, and *Ant4* still demonstrated CsA-sensitive mPT at increased matrix Ca^2+^ overload, and only deletion of the *Ppif* gene encoding CypD in these mice completely prevented Ca^2+^-induced mPT^26^. These findings suggested a new mPTP model consisting of two distinct molecular components, ANT and another CypD-regulated channel, that mediate mPTP formation^26^.

ATP synthase is a multi-subunit complex consisting of the F_1_ and F_O_ domains^27^ connected through the central and peripheral stalk subunits. The hydrophilic F_1_ domain is situated in the matrix, while most of the F_O_ subunits, including the c-ring, are hydrophobic, membrane-embedded proteins^27-29^. Conformational changes in the F_1_ domain that can be transmitted to F_O_ through the peripheral stalk were proposed to be crucial in activating the ATP synthase c-subunit leak channel (ACLC)^2, 30, 31^. Thus, compounds that specifically bind to ATP synthase may modulate its catalytic^32, 33^ (ATP synthesis and hydrolysis) and/or leak channel activities^17, 34^.

Bedaquiline (BDQ), also known as Sirturo, binds at the interface of F_O_ subunits a and c, inhibiting the catalytic activity of *Mycobacterium tuberculosis* ATP synthase^16^. It is also the first anti-mycobacterial drug approved by the Food and Drug Administration for treating pulmonary multidrug-resistant tuberculosis (TB)^35, 36^.

Prior studies have reported BDQ as a selective and potent inhibitor of mycobacterial, but not mammalian, ATP synthase^37^. However, BDQ was recently shown to bind and inhibit the catalytic activities of yeast and human ATP synthase^38, 39^.

Here, we report the new role of BDQ as a modulator of the mammalian ATP synthase c-subunit leak channel activity. We show that BDQ inhibits the single-channel activity of ACLC in a dose-dependent manner in planar lipid bilayer recordings of porcine heart ATP synthase. Additionally, we demonstrate that BDQ prevents glutamate-induced cell death in primary hippocampal neurons, indicating a vital role of ACLC in regulating cell death under excitotoxic conditions. These findings suggest the repurposed application of BDQ in treating ACLC-induced pathologies via inhibition of the ATP synthase leak channel activity.

## Results

### BDQ modulates the opening of the mitochondrial permeability transition pore

To study the effect of BDQ on the mPTP, a calcium retention capacity (CRC) assay was performed on isolated mitochondria from HEK293 cells. During the CRC experiment, isolated mitochondria were exposed to Ca^2+^ pulses to induce mPTP activation. Ca^2+^ accumulates in mitochondria until it triggers mPTP opening, followed by membrane depolarization, rupture of mitochondria, and release of sequestered Ca^2+^. The mitochondrial calcium uniporter (MCU) is the primary transporter facilitating Ca^2+^ uptake under these conditions. MCU-mediated Ca^2+^ transport depends on the mitochondrial membrane potential (ΔΨm), implying that only energized mitochondria can support its activity^40^. Succinate, a Complex II substrate, was used to energize the mitochondria. Fig. 1a-b shows that BDQ delayed the mPTP in a dose-dependent manner. BDQ significantly delayed the mPTP at higher concentrations and had a subtle effect at lower concentrations (Fig. 1a, b). ATP and CsA were used in this assay as positive controls. Mitochondria treated with these compounds demonstrated significantly delayed mPTP opening (Fig. 1a, b). These results suggest that BDQ delays mPTP opening by possibly preventing ACLC activation.

**Figure 1.**
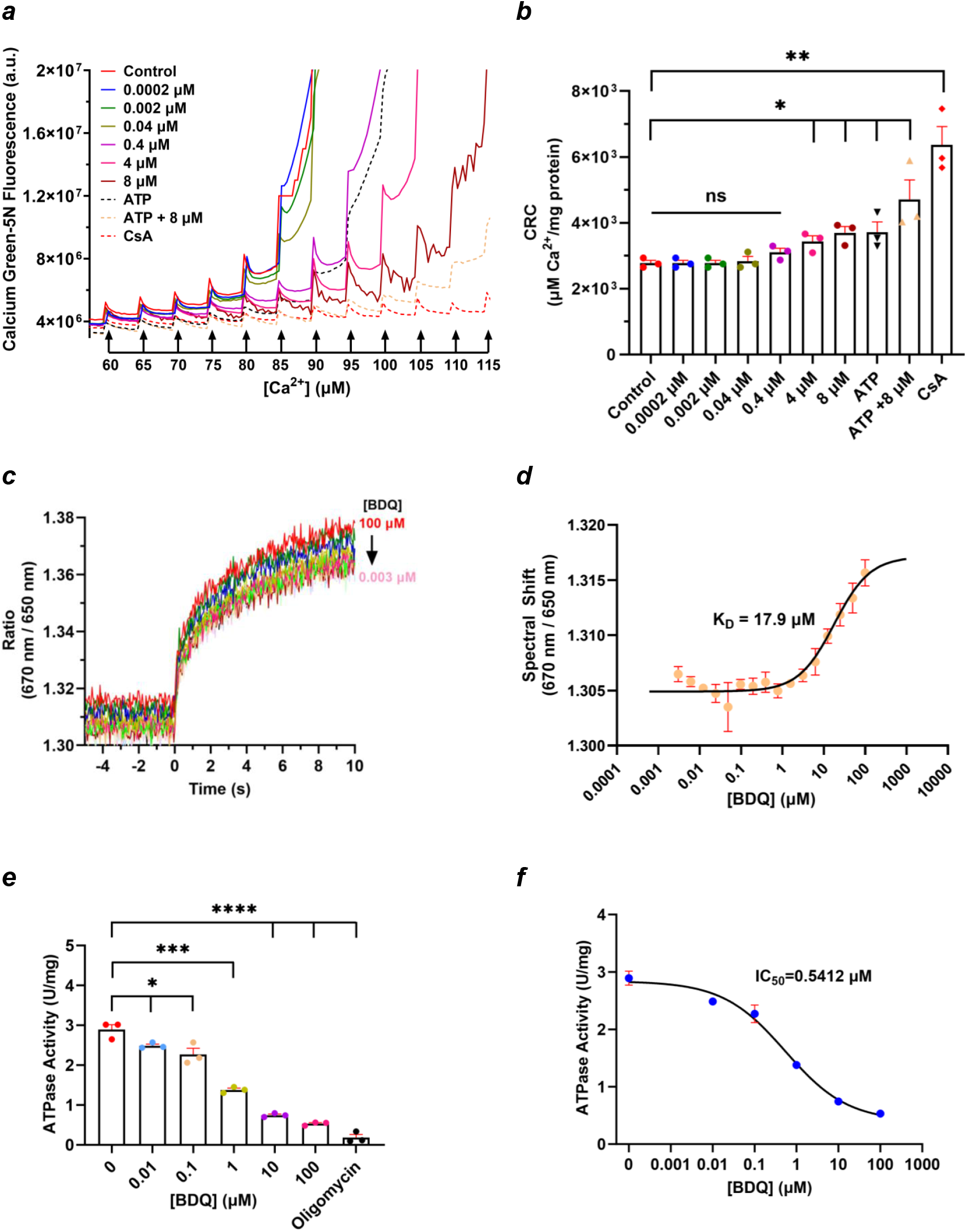
BDQ directly binds to porcine heart ATP synthase and inhibits its ATP hydrolysis activity. a. Mitochondrial calcium retention capacity (CRC) assay performed with isolated mitochondria in the presence of BDQ. **a**. Representative CRC assay in the presence of different concentrations of BDQ. Mitochondria (25 µg) isolated from HEK293 cells were used. ATP (400 µM) and CsA (10 µM) were used as positive controls. **b**. The group data for the calcium retention capacity assay (n=3). Error bars are represented as the standard error mean (SEM). The unpaired t-test was used for statistical analysis. 0.0002, and 0.002, μM BDQ, P >0.999; 0.04 μM BDQ, P=0.7268; 0.4 μM BDQ, P=0.0935; 4 μM BDQ, *P=0.0270; 8 μM BDQ, *P=0.0119; ATP *P=0.0427; ATP + 8 μM BDQ, *P=0.0326; CsA, **P=0.0030. **c**. Raw traces of a fluorescence signal of labeled ATP synthase in the presence of BDQ (100 µM-3 nM). The traces represent the ratio of signals measured at 670 nm and 650 nm. Samples were excited at t=0s, and spectral shifts were calculated over 10s. **d**. A dose-response curve was obtained from the spectral shift assay in the presence of different concentrations of BDQ. Data were fitted using MO Control 2 analysis software (NanoTemper Technologies). The presented data points are averages of 4 independent experiments, and the displayed error bars represent standard deviation (SD). **e**. ATP hydrolysis activity of porcine heart ATP synthase (4 µg) in the presence of BDQ (0.01-100 µM). Oligomycin (5 µg/µl) was used as a positive control (n=3). Error bars are represented as SEM. The unpaired t-test was used for statistical analysis. 0.01μM BDQ, *P=0.034; 0.1 μM BDQ, P=0.0338; 1 μM BDQ, *** P=0.0003; 10 μM BDQ, ****P<0.0001; 100 μM BDQ, ****P<0.0001; 5μg/μl oligomycin, **** P<0.0001. **f**. The fitted dose-response curve of the data is shown in **e**.

### BDQ inhibits the hydrolytic activity of mammalian ATP synthase

We studied the effect of BDQ on the catalytic and leak channel activities of porcine heart ATP synthase.

First, we measured the binding affinity of BDQ to purified ATP synthase using a spectral shift assay. The purified porcine heart ATP synthase was labeled with NHS-Red and titrated with a range of BDQ concentrations from 100 µM-3 nM. The raw values of the observed spectral shift for all samples were plotted and fitted, yielding a K_D_ value of 17.9 µM (Fig. 1c, d). The obtained binding affinity correlates with results observed for human ATP synthase, measured by surface plasmon resonance (K_D_ = 5.7 µM)^39^.

Different concentrations of BDQ (0.01 µM to 100 µM) were used to measure its effect on the ATP hydrolysis activity of purified porcine heart ATP synthase (Fig. 1e). The IC_50_ of BDQ for ATP hydrolysis inhibition was found to be ∼0.5 µM (Fig. 1f). Oligomycin was used as a positive control, and it completely inhibited the ATP synthase hydrolysis activity in this assay (Fig. 1e).

BDQ reported to bind to *M. tuberculosis*^*39*,41^, yeast^38^ and human ATP synthase^39^ at the interface of a-subunit and c-ring of F_O_ domain in the recent cryo-EM studies. In addition, NMR and mutagenesis studies show that BDQ can also bind to the ATP synthase F_1_ ε-subunit^42,43, 44^ with a K_D_ of 19.1 μM^44^. However, this binding site was not reported in any cryo-EM studies. Interestingly, the reported K_D_ closely matches the one found in our studies (17.9 μM, Fig. 1d), suggesting the possible binding of BDQ to either or both of these sites. Changes in the interactions in the F_1_ central stalk subunits (γ, δ, ε) with the F_O_ c-ring were suggested to play an important role in ACLC gating^24, 30^. Thus, conformational changes in ATP synthase induced either through binding of BDQ to ε-subunit or at the a/c-subunit interface may potentially modulate the leak channel activity of ATP synthase.

### BDQ inhibits the leak channel activity of mammalian ATP synthase

The effect of BDQ on the single-channel activity of purified porcine heart ATP synthase was studied in planar lipid bilayer recordings. Fig. 2a shows the dose-dependent inhibition of the ATP synthase leak channel by BDQ during a continuous channel recording. The first three additions of BDQ (0.01-0.03 µM) reduced the peak current from 150 pA to 82 pA at +50 mV, with further additions completely inhibiting the channel (Fig. 2a). Fig. 2b depicts a representative continuous recording of ACLC before and after the addition of 25 µM BDQ, showing complete inhibition of the channel. Similarly, lower concentrations of BDQ (1.5 µM and 6 µM) inhibited the channel in Fig. 2c and 2d. Fig. 2e shows no effect on the channel activity upon the first addition of BDQ (0.005 µM), while the subsequent additions (0.005 µM and 0.02 µM) induced complete closure of the channel. Fig. 2f represents the voltage ramp recording of ACLC in the presence and absence of bedaquiline at voltage values ranging from -100 mV to +100 mV. In this experiment, BDQ inhibits channel conductance at all voltages. Fig. 2g displays the group data of peak conductances in the presence of different concentrations of BDQ. The peak conductance of ACLC decreased gradually with increasing BDQ concentration (Fig. 2g). Fig. 2h shows the nonlinear fit of the data in Fig. 2g, suggesting that BDQ inhibits ACLC activity with an IC50 of 0.024 µM.

**Figure 2.**
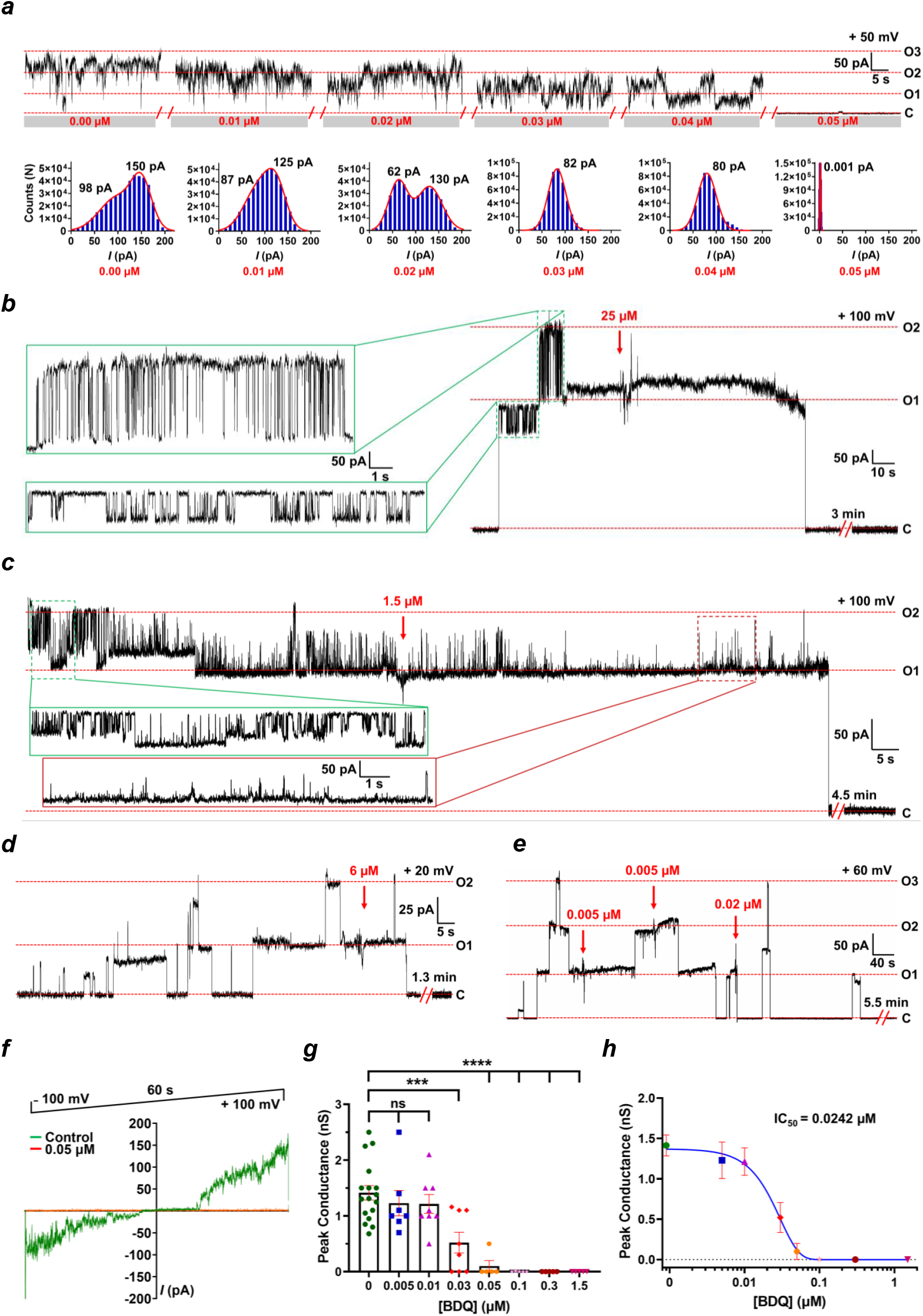
BDQ inhibits the porcine heart ATP synthase leak channel activity. **a**. A dose-dependent effect of BDQ on channel activity (0.01-0.05 µM). The truncated traces from a continuous planar lipid bilayer recording are displayed. BDQ was added at the *cis* side of the cuvette in 0.01μM increments. The lower panel is a histogram representation of the current, *I* (pA). The Gaussian fits are represented with a red outline, and the respective fitted peak current values are indicated. **b. c**. and **d**. A complete inhibition of the ATP synthase leak channel activity in the presence of 25, 1.5, and 6 µM BDQ, respectively. **e**. A channel inhibition upon the addition of three sequential doses of BDQ, as indicated by red arrows. **f**. Voltage-ramp recordings of the ATP synthase leak channel in the presence and absence of BDQ (0.05 µM). **g**. Group data on peak conductances in the presence of different concentrations of BDQ. **h**. The IC_50_ curve for BDQ inhibition based on the peak conductance values shown in **g**. The traces in the green boxes (panels **b**. and **c**.) display the channel activity before the addition of BDQ. The trace in the red box (panel **c**.) represents the channel activity after BDQ addition. “C” refers to the channel closed state, and "O" refers to the open state. Signals were filtered at 5 kHz using the amplifier circuitry. The error bars in **g**. and **h**. are represented as standard error mean (SEM) values. 0.005 μM BDQ (n=7), P = 0.4655; 0.01 μM BDQ (n=8), P = 0.3763; 0.03 μM BDQ (n=8), ***P = 0.0007; 0.05; 0.1; 0.3; 1.5 μM BDQ (n=5, for each concentration), ****P <0.0001.

### BDQ prevents glutamate-induced cell death in primary neuronal cultures

The excessive release of glutamate occurs in the brain during ischemia-reperfusion (I/R) injury^45^. To mimic I/R *ex vivo*, we treated primary hippocampal neurons with glutamate to measure the cell viability under excitotoxic conditions. The effect of BDQ on modulating glutamate-induced cell death was assessed by concurrently treating neurons with glutamate and different concentrations of BDQ. The well-known mPTP inhibitor, cyclosporin A (CsA), was shown to inhibit glutamate-induced cell death, suggesting that mPTP is a key cell death pathway under these conditions ^24^. We first evaluated the effect of varying concentrations (0, 0.1, 0.5, 1, and 5 µM) of BDQ under glutamate excitotoxic conditions (20 µM) to test its neuroprotective properties by measuring lactate dehydrogenase (LDH) release (Fig. 3a). Primary hippocampal neurons showed a dose-dependent response to BDQ treatment. In particular, treatment with 0.1 µM BDQ significantly protected neurons against excitotoxicity, whereas treatment with 0.5 or 1 µM BDQ did not show a significant neuroprotective effect. In contrast, 5 µM BDQ treatment exacerbated cytotoxicity (Fig. 3a).

**Figure 3.**
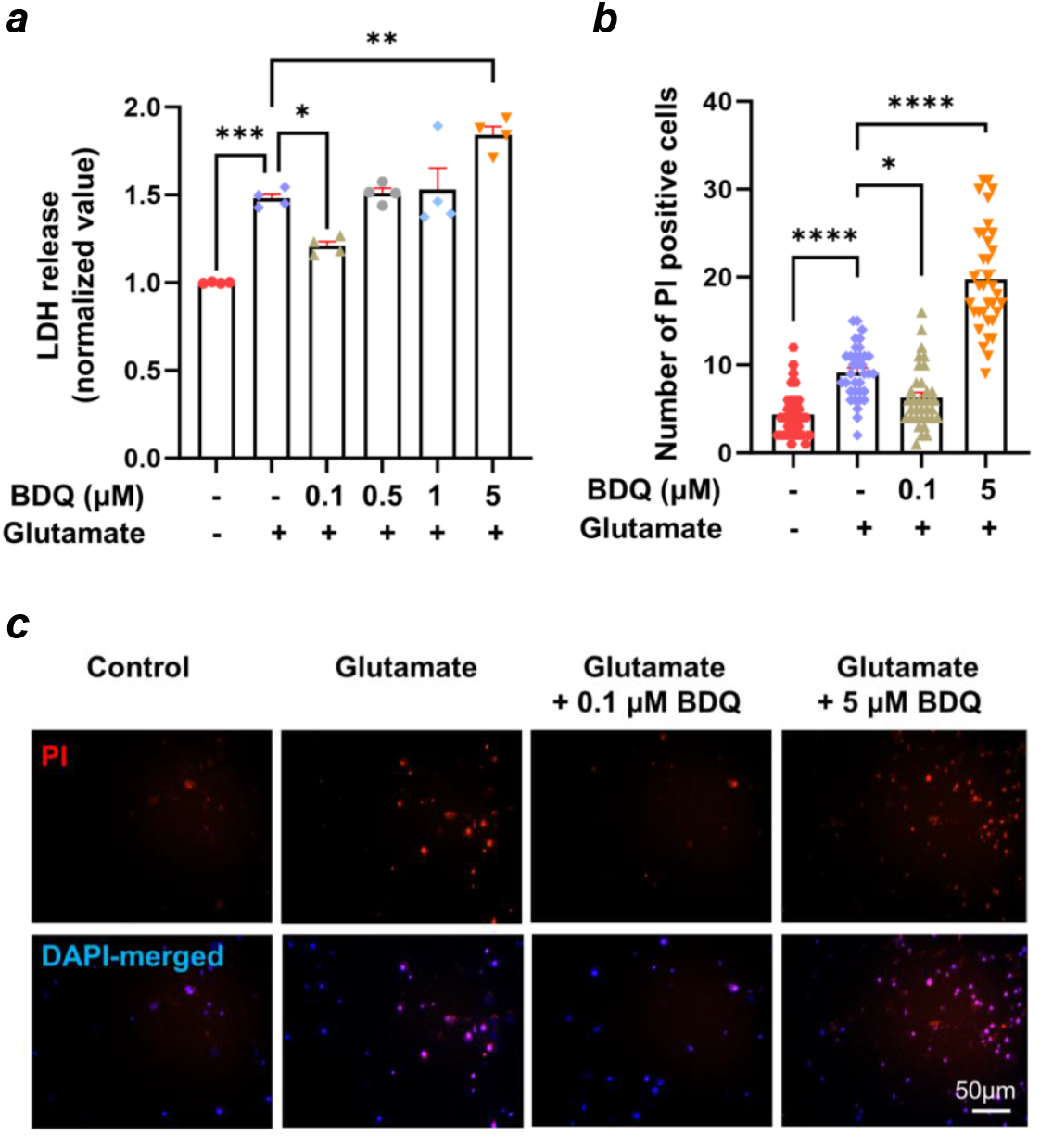
BDQ prevents glutamate-induced cell death in primary hippocampal neurons. **a**. The group data for lactate dehydrogenase (LDH) release, an indicator of cytotoxicity, of hippocampal neurons under the indicated conditions: Control, Glutamate (20μM), and BDQ (0.1, 0.5, 1, and 5 μM) with glutamate. Neurons were treated with Glutamate and Glutamate with BDQ for 24 h before any downstream assays (n=4); One-way ANOVA with a Tukey post hoc analysis was used. * P < 0.05, ** P < 0.01, and *** P < 0.001. **b**. Group data on propidium iodide (PI)-positive dead cells in cultured hippocampal neurons under the indicated conditions. n=35 micrographs from 3-5 biological replicates. A one-way ANOVA with Tukey post hoc analysis was used. * P < 0.05 and **** P < 0.0001, **c**. images of PI staining of cultured primary hippocampal neurons. Red: PI. Blue: DAPI. Scale bar = 50 μm.

We further quantified the cell death of glutamate-treated neurons by propidium iodide (PI) staining (Fig. 3b-c). The primary hippocampal neurons were treated with two different concentrations of BDQ (0.1 μM and 5 μM) along with the glutamate treatment. The low concentration of BDQ (0.1 μM) rescued cells from mPTP-induced cell death, confirming the role of BDQ as a potent inhibitor of mPTP via possible inactivation of ACLC under these conditions (Fig. 3a-c). Higher concentrations of BDQ, however, aggravated glutamate-induced cell death, most likely due to complete inhibition of the catalytic activity of ATP synthase.

## Discussion

Bedaquiline, a first-in-class diaryquinoline antituberculosis drug, prevents the growth and proliferation of *M. tuberculosis* by inhibiting the catalytic activity of ATP synthase to synthesize ATP^46, 47^. BDQ was reported to bind at the interface of the c-ring and the a-subunit of mammalian^39^ and *M. tuberculosis*^39, 48^ ATP synthase in cryo-electron microscopy structures.

In this study, we report a novel role for BDQ as a potent inhibitor of the mammalian ATP synthase leak channel. BDQ inhibits the single-channel activity of ACLC in a dose-dependent manner in planar lipid bilayer recordings. BDQ binds directly to porcine heart ATP synthase with a K_D_ of 17.9 µM as measured in the spectral shift assay. Additionally, BDQ treatment prevents glutamate-induced cell death in primary hippocampal neurons, suggesting an essential role for ACLC in neuronal death under these conditions and its possible role in mPTP. Nevertheless, we cannot rule out the potential effect of BDQ on the other mitochondrial proteins that may be involved in mPTP formation or its regulation.

mPTP is a voltage-gated, Ca^2+^- and Cyclophilin D (CypD)-regulated, large conductance leak channel situated in the mitochondrial inner membrane^49, 50^. The activation of mPTP, often triggered by mitochondrial Ca^2+^ overload and oxidative stress, leads to increased permeability of the mitochondrial inner membrane, mitochondrial swelling, and rupture, resulting in cell death^2, 3, 51^. mPTP is a non-selective leak channel that allows solutes up to 1.5 kDa to pass through the inner membrane^50, 52, 53^. It was reported to fluctuate between different open states (low- and high-conductance conformations) in mitoplast electrophysiological recordings^51, 52, 54-56^. The low-conductance flickering channel, with a mean conductance of ∼0.3 nS, contributes to the physiological function of mPTP^23, 56-58^. The high-conductance channel, with ∼1.5 nS peak conductance activity, leads to prolonged, irreversible openings, inducing inner membrane depolarization and cell death^2, 49, 59^. Although the exact molecular identity of mPTP is still debated, the mitochondrial ATP synthase c-ring remains one of the pore-forming candidates for this enigmatic channel^13, 14, 17, 18, 24, 60, 61^. Detailed information about the identity, structure, and regulation of mPTP is key in designing therapeutics to target its activity specifically.

BDQ preconditioning was reported earlier to mitigate ischemia/reperfusion-induced brain injury in Wistar rats after middle cerebral artery occlusion-reperfusion^62^. Intraperitoneal (i.p.) injection of BDQ (2 mg/kg) to male rats 60 min before transient occlusion of the middle cerebral artery has been reported to reduce infarct volumes and neurological deficits in rats^62^. While this study did not measure brain levels of BDQ, its peak concentration (C_max_) in the brain was estimated to be 0.02 μM based on another in vivo study^63^. This neuroprotective concentration of BDQ is very close to the IC_50_ of 0.024 μM for inhibiting the ACLC in planar lipid bilayer recordings (Fig.2h). It is also near the concentration of BDQ (0.1 μM) that protected hippocampal neurons from glutamate-induced cell death (Fig. 3), suggesting an important role of ACLC in preventing neuronal death in I/R.

In our studies, a higher concentration of BDQ (5 μM) aggravated glutamate-induced neuronal death, which can be explained by the full inhibition of ATP synthase catalytic activity. BDQ inhibits the ACLC activity with an IC_50_ of 0.024 μM (Fig. 2h) and ATPase activity with an IC_50_ of 0.54 μM (Fig. 1e, f). The lower concentration of BDQ (0.1 μM) used in this study rescued glutamate-induced neuronal death by inhibiting ACLC activity while allowing normal ATP synthase function. The 0.1 μM BDQ inhibits ATP synthase catalytic activity by only ∼20% (Fig. 1f). In contrast, a higher concentration of BDQ (5 μM) inhibits ATP hydrolysis activity by ∼75%. Although it also inhibits ATP synthase leak channel activity, it exacerbated glutamate-induced cell death in the neuronal viability study (Fig. 3). Thus, the cytotoxic effect of the higher BDQ concentration can be attributed to its ability to inhibit the catalytic activity of ATP synthase. This also implies that the BDQ is more selective in inhibiting the leak channel than in inhibiting ATP synthase catalytic activity at lower concentrations. Different IC_50_ values for ATP hydrolysis and ACLC inhibition may also reflect distinct binding sites of BDQ on ATP synthase^39, 42^.

We have previously shown that overexpression of human ATP synthase c-subunit aggravates glutamate-induced neuronal death in primary hippocampal neurons and H_2_O_2_-induced cell death in HEK293 cells^24^. In contrast, c-subunit knockdown (KD) in neurons undergoing glutamate excitotoxicity prevented cell death^24^. Here, we show that pharmacological inhibition of the ATP synthase c-subunit leak channel by BDQ mitigates glutamate-induced neuronal death. These data suggest that BDQ may prevent Ca^2+^-induced conformational changes within ATP synthase under glutamate excitotoxic conditions that otherwise lead to channel activation, depolarization of mitochondrial membrane potential, and cell death.

We recently suggested that non-reversible dissociation of ATP synthase F_1_ from the F_O_ domains occurs during long-lasting openings of ACLC in glutamate excitotoxic conditions, which are accompanied by repeated and profound inner membrane depolarization^24^. This marks the point of no return for the cell, since prolonged pore opening triggers outer membrane rupture and the release of cytochrome c, leading to the activation of downstream cell death pathways in detrimental pathological conditions (such as brain or heart ischemia or neurodegenerative diseases)^24^.

Our proposed model highlights the crucial role of the c-ring as the main pore-forming component of ACLC^14, 17, 24, 64^, and the roles of the F_1_ and e-subunit as the two gates of ACLC, which tightly regulate channel activity. According to this model, the binding of Ca^2+^ to the ATP synthase β subunits triggers conformational changes in the F_1_ domain that are transmitted to the central and peripheral stalk subunits, thereby activating the channel within the c-ring^24, 30, 31, 61^ (Fig. 4). The binding of BDQ at the interface between the c-ring and a-subunit may stabilize interactions between them, prevent structural changes within the ATP synthase that activate the channel, mitigating ACLC-induced cell death (Fig. 4).

**Figure 4.**
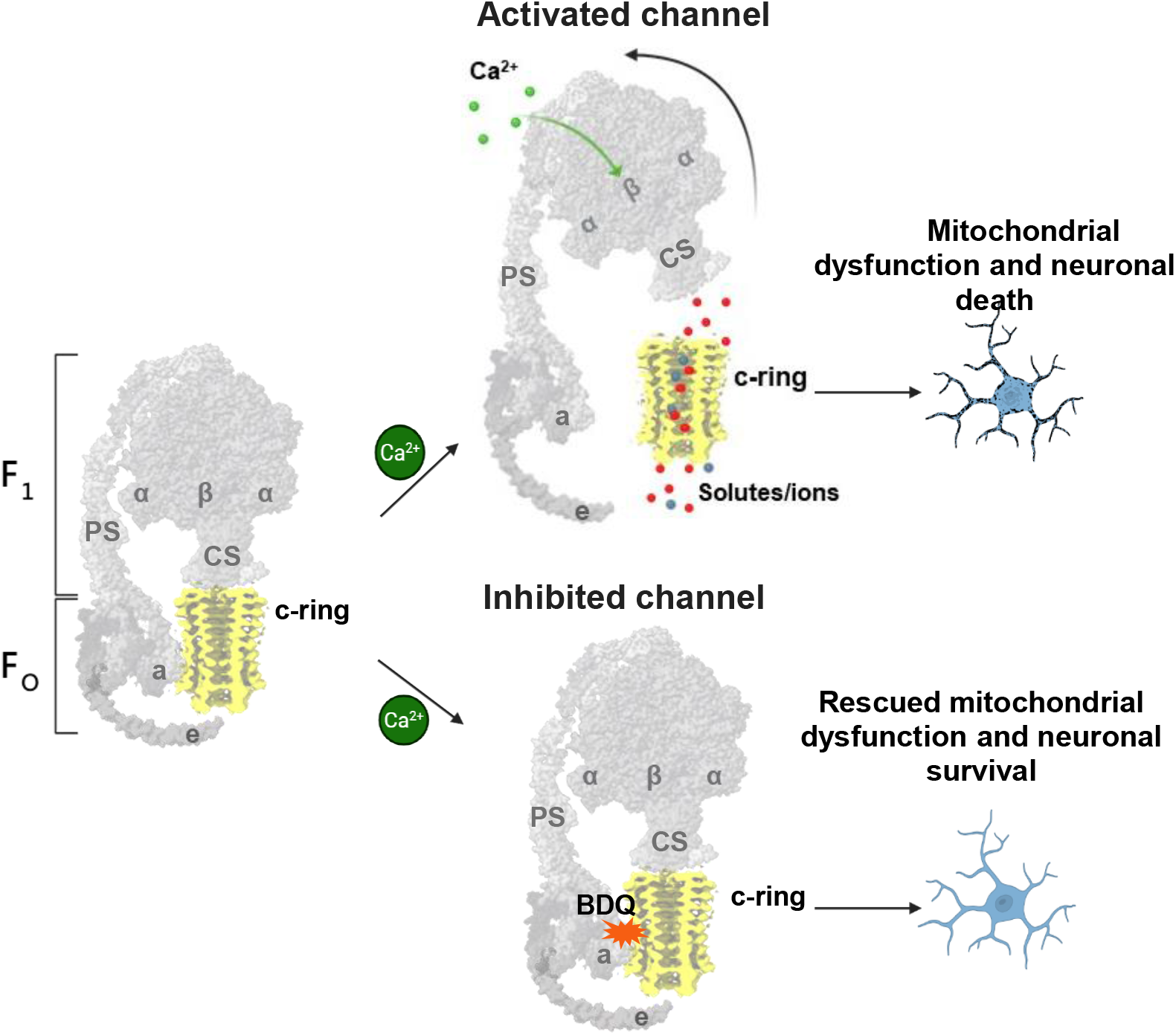
The proposed model of ATP synthase leak channel activation and the role of BDQ in its inhibition. The binding of Ca^2+^ to β-subunits may induce conformational changes in F_1_ that are transmitted to the central stalk (CS) and the peripheral stalk (PS) subunits, thereby activating the channel. During channel activation, the two gates of ACLC, F_1_ and the e-subunit, may move away from the c-ring, allowing channel formation and ion flow through the c-ring lumen. The binding of BDQ at the interface between the c-ring and a-subunit may stabilize interactions between them, prevent structural changes within the ATP synthase that activate the channel, and mitigate ACLC-induced cell death. The figure is created with BioRender.com. For simplicity, only the ATP synthase monomer is shown.

Our findings, supported by binding affinity, electrophysiological, and cell viability studies, reveal a novel role for BDQ as a specific inhibitor of the ATP synthase c-subunit leak channel. This study suggests the potential repurposing of this FDA-approved drug for the treatment of mPTP-related diseases.

## Materials and Methods

### Isolation of porcine heart mitochondria and purification of ATP synthase

Porcine heart mitochondria were isolated as described previously^17^. Briefly, the freshly collected porcine heart was finely minced on ice and transferred to 200 mL Buffer A (225 mM mannitol, 75 mM sucrose, 0.5 mM EGTA, 0.5% BSA, and 20 mM Tris; pH 7.4). Heart tissue was homogenized and centrifuged at 1000g for 10 min at 4^°^C. The supernatant was centrifuged at 8000 rpm for 15 min at 4^°^C. The obtained mitochondrial pellets were stored at -80^°^C for further use. ATP synthase was purified using differential centrifugation and size-exclusion chromatography as previously described^17^. Soluble ATP synthase fraction (in 0.5% DDM) was loaded into the Superose 6 column (Cytiva) equilibrated with 1.5 column volumes of elution buffer (20 mM Tris, 100 mM NaCl, 0.05% DDM, pH 7.8). Protein elution was performed in the same buffer, and 0.5 mL fractions were collected using an automated fraction collector (Cytiva). Fractions containing ATP synthase were pooled and used in experiments.

### Calcium retention capacity assay

Mitochondria were isolated from HEK293 cells using the Qproteome Mitochondrial Isolation Kit (Qiagen) according to the manufacturer”s protocol. Calcium Retention Capacity (CRC) was done with 25 µg of isolated mitochondrial protein in CRC buffer (150 mM Sucrose; 1 mM KH_2_PO_4_; 1 mM MOPS; 5 mM Succinate; 10 µM EGTA; pH 7.4) containing 1 µM Calcium Green-5N. 400 µM ATP and/or 10 µM CsA were used as a control. Fresh BDQ stocks were made in DMSO. The following concentrations of BDQ were used (8, 4, 0.4, 0.04, 0.002, 0.0002 μM/μg mitochondrial protein). Data acquisition was done with a microplate reader (SpectraMax− iD3; Molecular Devices) at an excitation and emission wavelengths of 485 and 535 nm, respectively. Calcium pulses (5 µM) were added every 2 min, and fluorescence data were recorded every 10s. The assay was done in a 96-well plate with a final volume of 100 µl. All the assays were performed at 23-25 °C.

### Spectral shift assay

Purified porcine heart ATP synthase was labeled with the Red-NHS 2nd-generation dye using a protein labeling kit (NanoTemper Technologies) according to the manufacturer”s protocol. Briefly, ATP synthase was incubated with Red-NHS dye, followed by removal of the free unreacted dye using the desalting column. Labeled ATP synthase (ATP synthase-Red-NHS) was used in the spectral shift experiments (Monolith X, NanoTemper Technologies). The experiment was performed at 10% excitation with medium IR laser power at 25^°^C. Spectral shift was measured for all the samples as a ratio of the fluorescence intensities at 670 nm and 650 nm, and is plotted against the logarithmic BDQ concentration. Sixteen serial dilutions of BDQ, ranging from 100 µM to 3 nM, were used in the experiments. The binding affinity, K_D,_ was determined from the obtained sigmoidal binding curve. Both the data acquisition and fitting were done using MO Control 2 software (NanoTemper Technologies).

### ATP hydrolysis assay of purified ATP synthase

ATP hydrolysis (ATPase) activity was measured in ATP cocktail buffer (50 mM Tris/H_2_SO_4_, 10 mM ATP, and 4 mM MgSO_4_, pH 8.0) at 37^°^C as described earlier^17^. One unit of enzymatic activity (U) corresponds to 1 μmol of ATP hydrolyzed (equivalent to 1 μmol of Pi produced) per min, per mg protein. The effect of BDQ on the ATP hydrolysis activity of porcine heart ATP synthase was determined in the presence of 0.01-100 µM BDQ. Inhibition activity data were fitted using the Dose-Response-Inhibition Model in GraphPad Prism (10.1.0) software. Oligomycin (5 µg/µl) was used as a positive control.

### Single-channel recordings of porcine heart ATP synthase

Planar lipid bilayer recordings were done in a 13 mm Derlin Cuvett (Warner Instruments) with an aperture of 200 µm. Lipids were prepared by drying a 15 µL aliquot of L-α-phosphatidylcholine (Avanti Polar Lipids; 20 mg/mL in chloroform) under N2 gas in a glass cuvette, then redissolving it in 60 µL of n-Decane (Thermo Scientific). After adding intracellular solution (120 mM KCl; 8 mM NaCl; 0.5 mM EGTA; 10 mM HEPES; pH 7.3) on both the *cis* and *trans* sides of the cuvette, the prepared lipid was directly painted onto the cuvette aperture using a fine paintbrush (Tintoretto, Italy). The stability of the bilayer was assessed by the resistance (5-7 GΩ) and C-pipette values (50-120 pF). Purified ATP synthase was added only on the *cis* side, and a constant voltage of ±40 to ±120 mV was applied to achieve protein insertion into the bilayer. A final concentration of 1 mM Calcium and 0.005 µM-25 µM bedaquiline was added on the *cis* side during the recordings to study its effect on the ATP synthase channel activity. Data acquisition was performed using an ePatch amplifier (Elements) with EZ Patch Software.

Clampfit software (Molecular Devices) was used for data analysis. The measured current was adjusted for the holding voltage, assuming a linear current-voltage relationship. The conductance (G) is expressed in nS, following the equation *G = I/V*, where *I* is in pA, and *V* is in mV. Group data were evaluated in terms of conductance, and the represented error bars are in ± SEM.

### Culture of primary hippocampal neurons

Primary rat hippocampal neurons were prepared from rat fetuses (Sprague-Dawley, day 18 of gestation; Envigo, Indianapolis, IN) as described previously^65-68^. Briefly, neurons (0.3 × 10^6^ cells/35mm plate) were seeded on poly-L-lysine-coated plates and grown in neurobasal medium supplemented with B-27, glutamine, and antibiotics (Invitrogen, Waltham, MA) for 20-22 days *in vitro* (DIV). *BDQ treatment:* Neurons were either treated with BDQ (0.1, 0.1, 1, and 5 µM, AdooQ Bioscience, Irvine, CA) or a vehicle (DMSO). BDQ was added to the hippocampal neurons 24 h before any downstream assays (PI staining, LDH assay) were performed. *Glutamate treatment:* Neurons were treated with 20 µM of glutamate (Sigma-Aldrich, St. Louis, MO) or vehicle for 24 h. Glutamate was freshly prepared in sterile PBS and added to the cell culture medium. The vehicle control for glutamate experiments was sterile PBS. All protocols were approved by the Institutional Animal Care Committee (IACUC) of the University of Alabama, Tuscaloosa, AL (23-11-7123).

### Viability assay

Lactate dehydrogenase (LDH) assay: The level of cytotoxicity in primary neurons was assayed by measuring leakage of LDH using an *in vitro* toxicology assay kit (Sigma-Aldrich, St. Louis, MO) as previously described^69^. In brief, the culture media and lysed cells were collected after 24 h of treatment. The LDH assay mixture was added to each sample. After a 20-minute incubation, the reaction was terminated by adding 1 N HCl. LDH activity was spectrophotometrically measured using a Clariostar microplate reader (BMG Labtech, Cary, NC) with an absorbance setting of 490 nm. Data were calculated by dividing the activity of LDH leaked into the medium from damaged cells by total LDH activity in the culture. *Propidium iodide (PI) staining:* Dead cells were stained with PI as previously described^14, 65, 68^. After treatment, 0.5 μM PI (Invitrogen) was added to the culture medium for 30 min at 37^°^C in the dark. Images were taken using a Zeiss AxioVert A1 microscope using a consistent exposure time. The number of PI-positive neurons was analyzed using AxioVision 4.9.

## Acknowledgments

We thank Dr. Evgeny Pavlov and Louis Betz for their insightful scientific discussions and constructive review of the manuscript. This work was supported by start-up funds from the Penn State College of Medicine and by a grant from the National Institute on Aging (R01AG072484) awarded to N. Mnatsakanyan.

## Author contributions

N.M. designed the project. A.K., E.S., I.M., S.E., Y.W, K.A.F., E.A., H.P. performed experiments and analyzed data. All authors contributed to the manuscript preparation.

## Declaration of interests

The authors declare no competing financial interests.

## Notes

### Competing Interest Statement

The authors have declared no competing interest.

